# Learning Dynamic Protein Representations at Scale with Distograms

**DOI:** 10.64898/2026.01.29.702509

**Authors:** Nicolas Portal, Wissam Karroucha, Vincent Mallet, Massimiliano Bonomi

**Author notes:** Correspondence to: Vincent Mallet < >, Massimiliano Bonomi < >.

## Abstract

Protein function and other biological properties often depend on structural dynamics, yet most machine-learning predictors rely on static representations. Physics-based molecular simulations can describe conformational variability but remain computationally prohibitive at scale. Generative models provide a more efficient alternative, though their ability to produce accurate conformational ensembles is still limited. In this work, we bypass expensive simulations by leveraging residue–residue distance probability distributions (distograms) from structure predictors such as Boltz-2. Our approach provides a scalable way to encode dynamic information into protein representations, aiming to improve function prediction without explicit conformational sampling. All code required to reproduce the experiments presented in this work is publicly available at https://github.com/anonymous-research-world/distodyn.

## 1. Introduction

Proteins perform a wide range of functions within cells. Recently, Machine Learning (ML) approaches have been developed to predict protein function, particularly their interactions with small molecules, RNA, DNA, and other macro-molecules. Some methods leverage information from a protein’s three-dimensional structure, while others rely on its amino acid sequence to take advantage of the more abundant sequence data. However, since structure encodes function more directly, structure-based methods generally outperform those based solely on sequence (Yan et al., 2023).

Structure alone is often not enough to understand protein functions. Biological systems populate a variety of conformational states, and their functions often emerge from the interplay between structural and dynamic properties. For example, conformational transitions (Nussinov et al., 2023), allosteric regulation (Wodak et al., 2019), and lig- and binding site flexibility (Alghamedy et al., 2018) play a crucial role to achieve specific functions. Experimental techniques such as nuclear magnetic resonance spectroscopy and cryo-electron microscopy provide valuable insights into biomolecular dynamics but remain limited in their ability to fully characterize complex conformational landscapes at atomistic resolution. Consequently, simulation-based and integrative approaches currently represent the most effective strategies to characterize protein dynamics (Hoff et al., 2024).

Simulation methods used to model protein conformational ensembles can be broadly categorized into physics-based approaches and generative models. The main limitation of physics-based methods, such as molecular dynamics (MD) simulations, lies in the prohibitive computational cost required to exhaustively sample complex conformational landscapes. Generative models are also not guaranteed to explore all relevant conformational states, and their accuracy ultimately depends on the quantity and quality of the data they have been trained on.

ML models aiming at predicting protein function ultimately need to account for the dynamic nature of proteins. While state-of-the-art ML approaches are not built to support multi-conformation inputs (Hu & Ohue, 2025; Cao et al., 2025), recent studies have explored the possibility to encode multiple conformations directly into protein representations. A recent approach aggregates different conformations into pairwise residue correlations, ultimately improving static representations (Guo et al., 2025). While promising, the method depends on the availability of extensive MD datasets.

In this work, we take a different approach, completely sidestepping the explicit generation of protein conformations. Specifically, we leverage the probability distributions over residue–residue pairwise distances (distograms) predicted by modern structure prediction methods such as AlphaFold2 (Jumper et al., 2021) and Boltz2 (Passaro et al., 2025). These distributions have recently been shown to capture prediction uncertainty as well as structural dynamics (Brotzakis et al., 2025; Schnapka et al., 2026; Sen et al., 2025; Savaş et al., 2025). Notably, distograms are obtained as byproducts of the structure prediction pipeline and are therefore orders of magnitude cheaper to compute than MD-derived correlation features (Figure 1).

**Figure 1.**
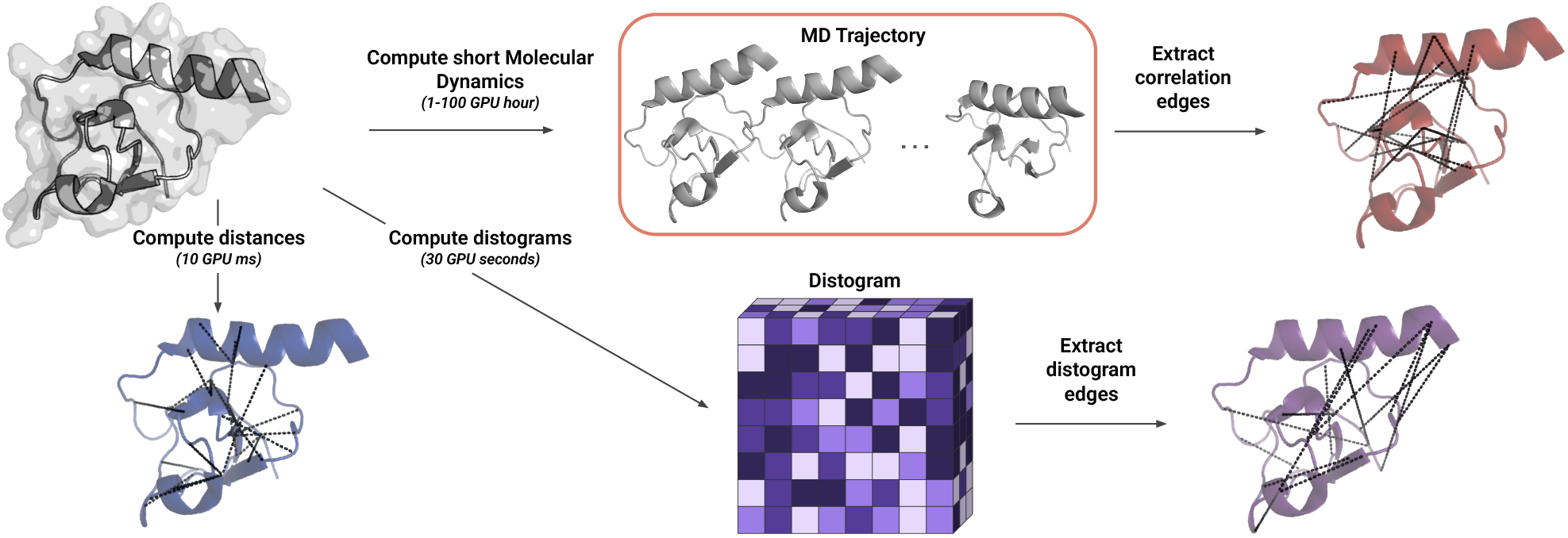
Overview of the protein representations used in this study. A protein can be represented as a graph extracted from a static structure. To capture dynamics, previous work relies on compute-intensive MD simulations (several GPU hours), encoding the results as additional edges. In contrast, we bypass simulations by leveraging predicted distograms (around 30 GPU seconds) to incorporate dynamic information.

**Figure 2.**
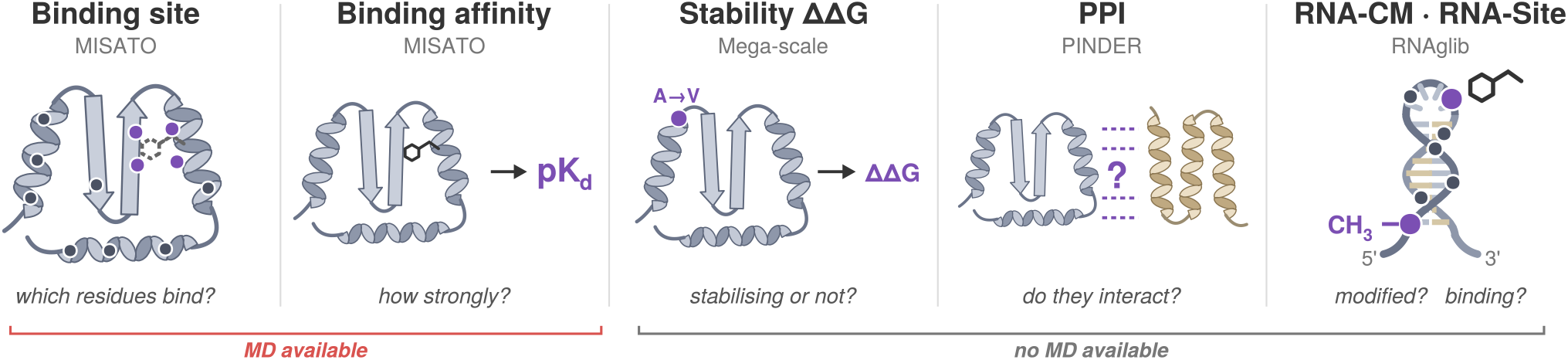
Visual overview of the different tasks we tested our method on. Predictors of binding site and binding affinity are trained on the MISATO dataset that gathers MD data, to be compared with other methods encoding dynamics (Section 5). Further validation on datasets where only static structures are available are presented in Section 6.

## 2. Related work

### Generating protein conformational ensembles

MD is currently the most widely used approach to generate protein and biomolecular conformational ensembles, drawing samples from the Boltzmann distribution under a given force field. Its high computational cost has motivated several strategies to improve sampling, including enhanced sampling techniques (Hénin et al., 2022), such as metadynamics (Laio & Parrinello, 2002). More recently, large-scale MD datasets have been released (Amaro, R. E. et al., 2025; Korlepara et al., 2024; Siebenmorgen et al., 2024; Vander Meersche et al., 2024), enabling generative models to be trained directly on MD data.

Generative models have consequently gained attention as a lower-cost alternative for generating conformational diversity (Klein et al., 2023; Schreiner et al., 2023; Jing et al., 2024; Wolf et al., 2025; Feng et al., 2025; Antoniadis et al., 2026), with some approaches approximating the Boltzmann distribution (Noé et al., 2019; Mardt et al., 2018; Lewis et al., 2025; Tan et al., 2025; Akhound-Sadegh et al., 2025; Lu et al., 2025; Roney et al., 2025; Kapusniak et al., 2026). Another class modifies modern structure prediction models to generate conformational diversity, for example by manipulating multiple sequence alignments (del Alamo et al., 2022; Wayment-Steele et al., 2024; Kalakoti & Wallner, 2025; 2026) or steering diffusion (Richman et al., 2025; Lam et al., 2026). Some of these models reproduce experimental observables, including NMR order parameters, small-angle X-ray scattering profiles, and temperature-dependent ensemble properties (Lewis et al., 2025; Janson et al., 2025).

### Protein representation learning

Geometric deep learning encoders have been widely used for protein representation learning (Isert et al., 2023), with representations based on 3D convolutions (Jiménez et al., 2017; Weiler et al., 2018), sequences (Rao et al., 2021), surfaces (Gainza et al., 2020), graphs (Aumentado-Armstrong, 2018), and equivariant discrete networks (Jing et al., 2021). Other approaches were designed specifically for protein structures, incorporating protein-specific properties into the network (Zhang et al., 2022; Hermosilla et al., 2020; Fan et al., 2022; Wang et al., 2022; 2025).

Multi-modal representations can combine complementary biological and computational priors. A common strategy combines sequence information with a structural graph (Hermosilla et al., 2020; Fan et al., 2022; Wu et al., 2023; Zhang et al., 2023), while other methods incorporate structural information into the training of protein language models (Bepler & Berger, 2019; Heinzinger et al., 2024; Su et al., 2023). More recently, methods combining multiple structural representations have also shown strong performance (Somnath et al., 2021; Zhang et al., 2024; Mallet et al., 2025).

Several methods incorporate MD simulations into ML-based representations. Some treat different conformations as data augmentation (Wu et al., 2022; Min et al., 2024; Libouban et al., 2025), while others use multi-instance learning (Ilse et al., 2018) to encode conformations independently and aggregate their representations (Zankov et al., 2021; Kleiman et al., 2025). The latter increases inference time for limited performance gains (Criscuolo et al., 2024). Finally, some methods aggregate conformations into a composite graph (Chiang et al., 2022; Kalifa et al., 2025; Guo et al., 2025). All these approaches rely on MD trajectory datasets.

## 3. Motivation and contributions

To encode dynamic information, Guo et al. (2025) proposed using residue-residue motion correlations extracted from MD trajectories as pairwise relationships to enrich the radius graph traditionally used in graph-based protein representation learning. Training relational graph neural networks on these enriched graphs improves performance across various tasks.

Here, we follow a similar strategy by extracting dynamic information from distograms. Distograms are distance distributions between *C*_*β*_ atoms (or *C*_*α*_ for glycines, *C*2 for RNA pyrimidines, and *C*4 for RNA purines). First introduced in methods presented at CAPRI13 (Senior et al., 2020; Xu & Wang, 2019), they are now provided by most state-of-the-art structure predictors, such as Boltz2, as equally spaced bins spanning 0.2 to 2.2 nm, with the last bin also capturing larger distances.

By representing the full probability distribution over internode distances, distograms capture both spatial proximity and structural uncertainty (Brotzakis et al., 2025; Schnapka et al., 2026; Sen et al., 2025; Savaş et al., 2025). Contrary to MD, they can be generated at scale for training and deployed at inference time.

In this paper:

- We enrich distance-based residue graphs with edges and features derived from distograms (Section 4).
- We show that these distogram-augmented graphs outperform graphs encoding MD trajectories (Section 5).
- We successfully apply our framework to protein stability, protein-protein interaction and RNA property tasks, where no MD data is available (Section 6).

Our work enables function prediction beyond static structures at limited computational cost.

## 4. Graph construction

Protein structures are represented by a graph *G*_**P**_ = (*V, ℰ* ). Depending on the task, *V* corresponds to either atoms or residues represented by their *C*_*α*_ atom. Node features are defined as one-hot encoding of the amino acid type or atom type. Traditionally, edges connect residues close in three-dimensional space, as determined by k-nearest neighbors or a radius cutoff. In this work, we used radius graph edges, defined as *ℰ* _*dist*_ = *{* (*v*_*i*_, *v*_*j*_) |*d*(*v*_*i*_, *v*_*j*_) *< τ*_*dist*_ *}*, where *d*(*v*_*i*_, *v*_*j*_) represents the distance between nodes *v*_*i*_ and *v*_*j*_, and *τ*_*dist*_ is a distance threshold.

When an MD trajectory is available, we follow the approach of Guo et al. (2025) and compute the correlation between the motions of residues *v*_*i*_ and *v*_*j*_, denoted as |*C*_*ij*_|. We define correlation-based edges *ℰ*_*corr*_ = *{* (*v*_*i*_, *v*_*j*_) ||*C*_*ij*_ *> τ*_*corr*_ *}*, where *τ*_*corr*_ is a correlation threshold. Following Guo et al. (2025), *τ*_*dist*_ and *τ*_*corr*_ are set to 10 Å and 0.3 respectively, (4.5 Å and 0.6 for atomic-graphs). Finally, Chroma’s developers (Ingraham et al., 2023) suggested adding random edges in protein graphs. These edges enable long-range message passing and represent an important negative control for our approach. We sampled edges uniformly, taking as many samples as the number of distogram edges, to obtain *ℰ*_*rand*_ *∼ U* (*V × V*), s.t. |*ℰ*_*rand*_| = |*ℰ*_*dist*_|.

### 4.1 Distogram-based edges and edge features

Distograms *D* encode the probability distribution of the distance between each pair of residues (*v*_*i*_, *v*_*j*_), effectively representing prediction uncertainty and, possibly, variability due to the underlying dynamics (Brotzakis et al., 2025; Schnapka et al., 2026; Sen et al., 2025; Savaş et al., 2025). In practice, these probabilities are discretized into *B* bins, (*b*_1_, *b*_2_, *·, b*_*B*_), with *b*_1_ = 0 and the convention that *b*_*B*+1_ =*∞* . Distograms are therefore tensors of shape (*N, N, B*), where *N* is the number of residues in the graph. The value at position (*i, j, k*) corresponds to the predicted probability that the distance between nodes *v*_*i*_ and *v*_*j*_ falls into bin *k*, i.e. *D*(*i, j, k*) = ℙ [*b*_*k*_ *≤ d*(*v*_*i*_, *v*_*j*_) *< b*_*k*+1_].

Based on these probabilities, we can define a distogram-based edge set composed of pairs predicted to be close with sufficient probability,

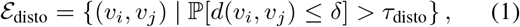

where *τ*_disto_ is a probability threshold and *δ* is a distance cutoff for the distogram-based neighborhood.

The value of *δ* is determined from the distribution of distances between neighboring residues as measured in the Protein Data Bank (PDB), and depends on the specific pairs (*u, v*) of amino-acid types as *δ*_*u,v*_ = *µ*_*u,v*_ + 1.645*σ*_*u,v*_ where *µ*_*u,v*_ and *σ*_*u,v*_ are tabulated for each amino-acid pair (Kamisetty et al., 2013). To be used in the discrete setting of distograms, *δ*_*u,v*_ is translated into a cut-off bin defined as the bin closest to the cutoff distance, 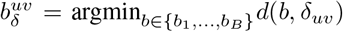. The probability in Eq. 1 is then computed as,

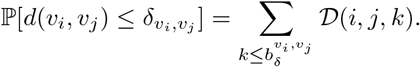

In our experiments, distograms were extracted from Boltz2 (Passaro et al., 2025) after softmax normalization. Our results were obtained with *τ*_disto_ = 10^*−*4^, which results in roughly the same number of edges as when computed from MD correlations. In addition to enriching graphs with additional edges, we can use the probability distributions *D*(*i, j*) *∈* Δ^*B*^ as edge features. This allows the model to capture both the expected distances and the dynamic variability between nodes.

### 4.2 Graph analysis

We analyze the different edge types introduced above using the MISATO dataset (Siebenmorgen et al., 2024), a collection of thousands of short MD simulations of protein-ligand complexes. A detailed description of this dataset is provided in Appendix A.3.1.

We first compute the intersection of the different edge types as a fraction of all possible pairs (Figure 3A). There is fair agreement between edge types: notably, distance edges (3.1% of possible pairs) are largely covered by distogram and correlation edges, with only 0.1% specific to distance. Correlation and distogram edges show moderate overlap.

**Figure 3.**
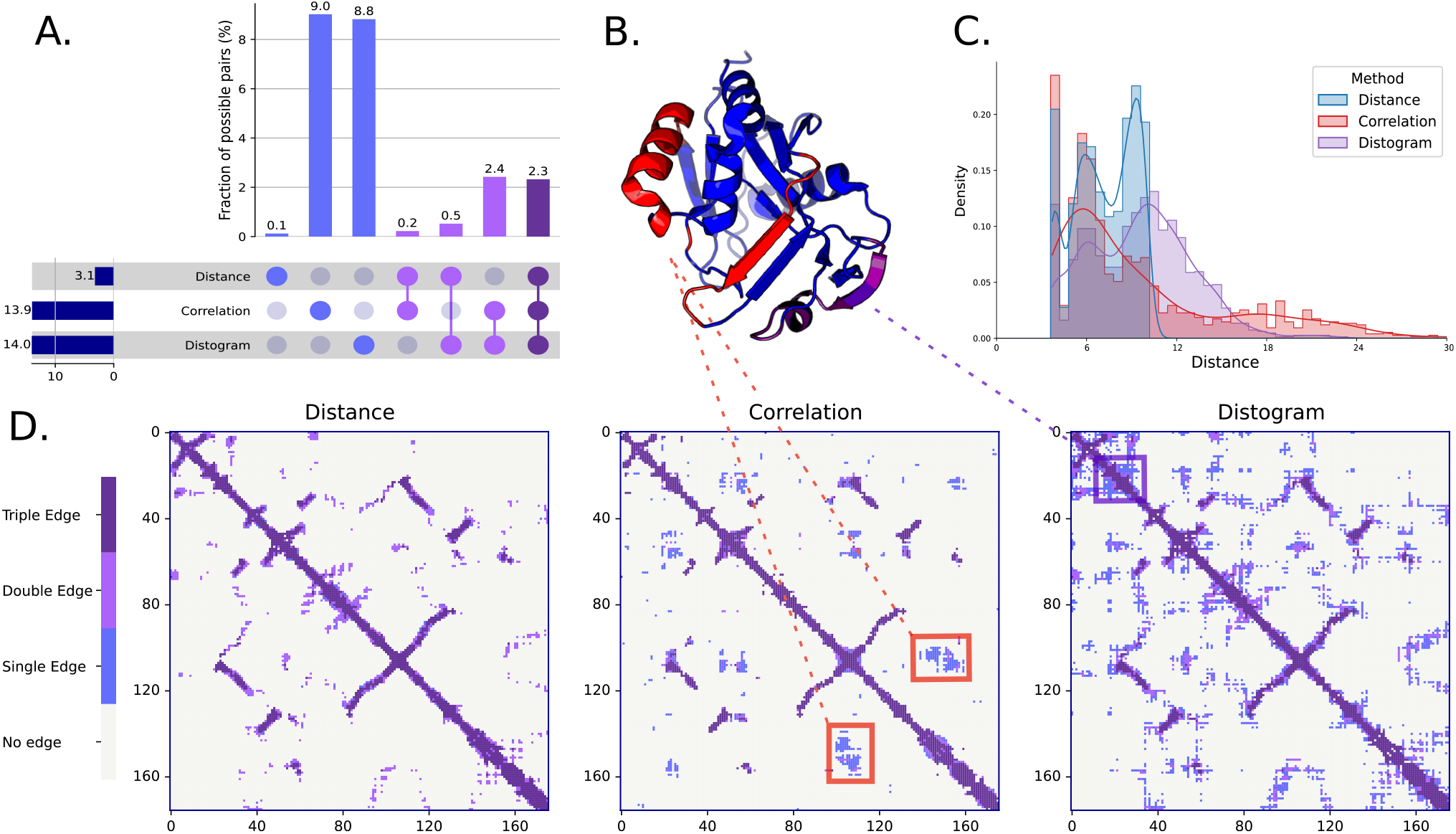
**A**. Upset plot showing the overlap between our edge types (*ℰ*_*dist*_, *ℰ*_*corr*_, and *ℰ*_*disto*_) across the binding site task test set. Rows represent individual edge types, while columns indicate specific combinations. **B**. 3D structure of the `6nvd` protein. Red and purple residues correspond to the red box in the correlation adjacency matrix and the purple box in the distogram adjacency matrix, respectively. **C**. Distance distributions for the different edge types in `6nvd`. **D**. Adjacency matrices for each edge type in `6nvd`. Colors indicate whether a residue pair appears in one, two, or all three edge types.

To illustrate their differences, we consider the 6nvd protein from our test set, an enzyme involved in biotin synthesis in the bacteria responsible for tuberculosis. Figure 3C shows the edge lengths for each type. The means are higher for *ℰ*_*corr*_ (9.63,Å ) and *ℰ*_*disto*_ (9.57,Å ) than for *ℰ*_*dist*_ (7.16,Å ). This trend is even stronger across the full dataset, with mean distances of 17.1,Å, 19.2,Å, and 7.2,Å, respectively. Thus, both correlation- and distogram-based edges enable long-distance message passing.

These edge types also exhibit distinct patterns (Figure 3D). Correlation-specific edges mapped to the structure (Figure 3B, red) correspond to a helix and a beta-sheet motif that move in a correlated manner. Thus, these edges need not represent proximity in alternative conformations. In contrast, distogram edges are found around distance edges, encoding structural fuzziness. We also identified distogram-specific edges (Figure 3B, purple) corresponding to a flexible loop with highly dynamic secondary structure.

### 4.3 Relational graph neural networks

To effectively leverage the diverse types of edge features described above (distance, correlation, and distogram-based), we employ *relational graph neural networks* (R-GNNs) that are designed to handle graphs with multiple relation types. In this framework, each edge type corresponds to a distinct relation, allowing the model to learn edge-specific message passing rules and therefore capture the different structural and dynamic properties encoded by each edge feature.

We experiment with three R-GNN variants: Relational Graph Convolutional Networks (R-GCN) (Schlichtkrull et al., 2018), Relational Graph Attention Networks (R-GAT) (Busbridge et al., 2019), and a Relational version of the Equivariant Graph Neural Network (R-EGNN) (Satorras et al., 2021). A detailed description of these models is provided in Appendix A.2. By using these relational GNN architectures, the model can capture the complementary information encoded in different edge types, while simultaneously leveraging node features to predict properties at the residue or atom level.

## 5. Validation on datasets where MD is available

We first evaluate our distogram-augmented relational graph architectures in scenarios where explicit dynamic data is available from short MD trajectories, benchmarking directly against correlation-based graphs (*ℰ*_*corr*_).

### Experimental setup

Following Guo et al. (2025), we evaluate on ligand binding site (443 nodes on average) and binding affinity (47 nodes on average) prediction. For each task, models are trained using distance, correlation, and distogram edges, both in isolation and combined as distinct relation types. Details about the dataset, learning and architecture hyperparameters are provided in Appendix A.3.1. Results for the RGAT models are presented in Table 1 and for other models in Appendix B.1.

**Table 1.** Performance of R-GAT on binding site and binding affinity prediction. We compare various dynamic-encoding approaches using *ℰ*_*dist*_ alone or combined with *ℰ*_*corr*_ or *ℰ*_*disto*_, optionally adding distogram-based edge features. Best-performing models are **bold**, second-best underlined.

| Graph Type | Binding Site Prediction |  |  |  |
| --- | --- | --- | --- | --- |
| | Accuracy (↑) | Precision (↑) | Recall (↑) | F1 (↑) |
| Distance | 0.753 | 0.196 | 0.478 | 0.278 |
| + Correlation | 0.791 | 0.229 | 0.469 | 0.308 |
| + Distogram | <u>0.814</u> | <u>0.289</u> | <u>0.598</u> | <u>0.390</u> |
| + Distogram + Features | **0.816** | **0.298** | **0.627** | **0.404** |

| Graph Type | Binding Affinity Prediction |  |  |  |
| --- | --- | --- | --- | --- |
| | MAE (↓) | RMSE (↓) | Pearson (↑) | Spearman (↑) |
| Distance | 1.226 | 1.562 | 0.691 | 0.658 |
| + Correlation | 1.280 | 1.568 | 0.686 | 0.656 |
| + Distogram | **1.176** | **1.487** | **0.740** | **0.713** |
| + Distogram + Features | <u>1.179</u> | <u>1.492</u> | <u>0.722</u> | <u>0.693</u> |

### Performance analysis

Incorporating distogram-based edges and features consistently and substantially improves performance, with distogram edges enhancing distance-based graph *across all models and metrics considered*.

We also compare our method to the correlation-based approach proposed by Guo et al. (2025), which relies on compute-intensive MD simulations. Our distogram-based approach clearly outperforms theirs, ranking first in 21/24 settings. Importantly, it remains competitive when ranking second, trailing only 2 accuracy points for R-GCN, but when ranking first gains can be substantial (e.g. 9.6 F1 points on binding site prediction).

Using distograms as explicit edge features yields mixed, architecture-dependent effects. For instance, edge features benefit R-GAT on binding site prediction but slightly hinder affinity estimation, whereas the inverse holds for R-EGNN. Since it sometimes results in strong improvements (Affinity, R-EGNN, +5 Spearman points), we advise users to test both approaches for their specific problem.

To investigate the information contained in correlation and distograms, we conducted two control experiments. First, we trained models on an empirical distogram constructed from the MD data and obtained results similar to the correlation approach. This direct comparison validates that our gains do not stem from methodological artifacts. Moreover, we tried to combine correlation edges with distogram edges, resulting in performance similar to using only distograms. We conclude that most of the information provided by short MD simulations is already encapsulated in our distograms. More details are provided in Supplementary section B.2.

Overall, our results indicate that distograms capture complementary structural information that is neither encoded by static distance thresholds nor correlation-based neighborhoods. This added information brings benefits to both classification on larger graphs and regression tasks on smaller graphs.

### Rewiring effects

The benefits of adding edges could correspond to either capturing neighborhoods in alternate conformations or to rewiring the graph by adding long-range edges. To disentangle these effects, we compared against two baselines. The first consists of adding random edges (Ingraham et al., 2023) which introduces unoptimized long-range connections like distogram edges. The second is a principled, topology-aware rewiring baseline (Topping et al., 2022) designed to avoid oversquashing. Table 2 highlights an illustrative case using R-GCN on binding site and affinity prediction. A full exploration across all models, and rewiring step counts is detailed in Appendix B.3.

**Table 2.** Rewiring results for the R-GCN model. Rewiring improves performance less than distogram edges, and both can be synergistically ombined.

| Graph Type | Binding Site Prediction |  |  |  |
| --- | --- | --- | --- | --- |
| | Accuracy (↑) | Precision (↑) | Recall (↑) | F1 (↑) |
| Distance | 0.761 | 0.187 | 0.420 | 0.259 |
| Distance + Distogram | 0.782 | <u>0.236</u> | <u>0.535</u> | <u>0.328</u> |
| Distance + Random | 0.772 | 0.191 | 0.401 | 0.259 |
| Distance + Rewired | <u>0.784</u> | 0.228 | 0.491 | 0.311 |
| Distance + Rewired + Distogram | **0.790** | **0.253** | **0.568** | **0.350** |

| Graph Type | Binding Affinity Prediction |  |  |  |
| --- | --- | --- | --- | --- |
| | MAE (↓) | RMSE (↓) | Pearson (↑) | Spearman (↑) |
| Distance | 1.244 | 1.562 | 0.689 | 0.656 |
| Distance + Distogram | <u>1.195</u> | <u>1.487</u> | <u>0.731</u> | <u>0.694</u> |
| Distance + Random | 1.368 | 1.715 | 0.649 | 0.619 |
| Distance + Rewired | 1.265 | 1.583 | 0.679 | 0.640 |
| Distance + Rewired + Distogram | **1.159** | **1.457** | **0.737** | **0.707** |

While extensive rewiring (1,000-10,000 loops) improves over random edges, distogram-induced edges still outperform it in most cases, confirming that the increase in performance is primarily driven by capturing alternate conformations. Notably, when combining rewired distance edges with distogram edges, we found that these two effects are synergistic, and even beneficial when rewiring alone was not. *Overall, the best performing method makes use of distograms across all models, metrics and tasks*.

## 6. Applications on generic datasets

In this section, we evaluate the performance of our approach in predicting various protein and RNA properties. In all tasks, dynamics play a key role. However, MD simulations datasets are not available which makes these tasks particularly challenging and highlights the need for accurate prediction methods that do not rely on such data.

### 6.1. Prediction of mutation effects

Protein stability is measured as the change in free energy Δ*G* between its folded and unfolded states. Dynamics in both folded and unfolded states plays an important role in determining protein stability (Frellsen et al., 2025). Upon mutations in the amino acid sequence, stability variations are quantified by ΔΔ*G*. Predicting those variations is crucial for protein design.

ThermoMPNN (Dieckhaus et al., 2024) was trained on the mega-scale dataset (Tsuboyama et al., 2023) to predict ΔΔ*G* from the native structure and mutated sequence. To extract structure embeddings, ThermoMPNN relies on a distance-based graph model (Dauparas et al., 2022), that we sought to enrich with distograms. Data and splits were held constant (more details in Appendix A.3). We also include results for SPURS (Li & Luo, 2025), a recent baseline tailored for this task.

Given the small size of the proteins studied here (56 nodes on average) and the high number of neighbors used by ProteinMPNN (48 nearest neighbors), little room is left for adding edges. Therefore, we only investigate adding edge features to the baseline model. We present the results of our experiments in Table 3.

**Table 3.** Protein stability results, computed over five seeds. Including distogram as edge features enhances ThermoMPNN results and obtains SOTA performances.

| Model | $R^2$ (↑) | RMSE (↓) | Spearman R (↑) | Pearson R (↑) |
| --- | --- | --- | --- | --- |
| Spurs (Li & Luo, 2025) | - | - | 0.74 | 0.78 |
| ThermoMPNN | $0.524 \pm 0.012$ | $0.722 \pm 0.009$ | $0.722 \pm 0.006$ | $0.755 \pm 0.006$ |
| + Distogram Features | $0.567 \pm 0.018$ | $0.689 \pm 0.014$ | $0.743 \pm 0.003$ | $0.782 \pm 0.005$ |

Our results show that incorporating distogram features clearly improves performance across all metrics. The *R*^2^ increases from 0.524 to 0.567, RMSE decreases from 0.722 to 0.689, and both Pearson and Spearman correlations improve. This indicates that distograms capture structural variability relevant for stability prediction, providing informative edge features that complement ProteinMPNN. SPURS improved over ThermoMPNN by coupling ProteinMPNN with ESM2 and a more elaborate architecture. Our approach reaches the performance reported for SPURS without architectural modifications.

### 6.2. Prediction of protein-protein interactions

The prediction of mutation effect tasks above involves comparatively small graphs. To test distogram-based edges on substantially larger protein graphs, we apply our approach to protein-protein interaction (PPI) prediction. The state-of-the-art predictor, SpatialPPIv2 (Hu & Ohue, 2025), relies on a graph representation. We add distogram edges and edge features without changing the architecture. Models are built on the large-scale PINDER dataset (Kovtun et al., 2024), that covers markedly larger graphs than in the task above (162 residues on average) and includes a structurally deleaked test set. We restrict our analysis to heterodimers, as almost all homodimers in PINDER represent positive pairs. We report additional details in Appendix A.3.3 and our results in Table 4.

**Table 4.** Results on protein heterodimers prediction, on the PINDER dataset. We enrich the SpatialPPIv2 model with distogram information and observe consistent gains.

| Method | Accuracy | Precision | Recall | F1 | AP | AUC-ROC |
| --- | --- | --- | --- | --- | --- | --- |
| SpatialPPIv2 | 0.777 | 0.916 | 0.609 | 0.732 | 0.900 | 0.916 |
| Distance + Distogram | 0.827 | 0.903 | **0.732** | 0.809 | 0.926 | 0.930 |
| Distance + Distogram + Features | **0.835** | **0.929** | 0.726 | **0.815** | **0.941** | **0.945** |

Adding distogram edges, and adding distogram edge features, consistently and substantially improves performance over the SpatialPPIv2 baseline across all metrics, most notably raising the F1 score from 0.732 to 0.815 and accuracy from 0.777 to 0.835. These results indicate that the benefits of distogram-based representations extend to larger protein graphs, and establish a new state of the art on this task.

### 6.3. Prediction of RNA properties

Finally, we adapt our approach to RNA molecules, typically more flexible than proteins. We use the RNA-CM and RNA-Site tasks, introduced in the RNAglib benchmark (Wyss et al., 2025). RNA nucleotides can be chemically modified, subtly altering their shape but crucially affecting their functions. The RNA-CM task aims to predict these chemical modifications from an RNA structure. The RNA-Site task is similar to the aforementioned ligand binding site prediction in proteins. These tasks have 57 and 64 nodes on average, respectively.

Considering the limited size of the RNA graphs, we only investigate the impact of adding distogram edge features to different graphs representing RNA: distance-based graph, 2D+ graph (encoding backbone and canonical base pairs), and 2.5D graph (additionally encoding noncanonical base pairs, as defined in Leontis & Westhof (1998)). We compare our edge features to distance encoded with an RBF. The results are presented in Table 5, and additional details are provided in Appendix A.3.4.

**Table 5.** Balanced accuracy results of different constructions on the RNA-CM and RNA-Site tasks. We report standard errors computed over three seeds.

| RNA-CM |  |  |  |
| --- | --- | --- | --- |
|  | Distance | 2D+ | 2.5D |
| None | **$59.6 \pm 1.3$** | $62.1 \pm 0.4$ | $59.8 \pm 1.3$ |
| Distance | $56.9 \pm 0.2$ | $58.2 \pm 0.5$ | $60.1 \pm 1.4$ |
| Distogram | $55.2 \pm 0.4$ | **$64.7 \pm 0.1$** | **$68.2 \pm 0.9$** |
| RNA-Site |  |  |  |
|  | Distance | 2D+ | 2.5D |
| None | $48.6 \pm 0.3$ | $57.7 \pm 1.1$ | $56.1 \pm 0.3$ |
| Distance | $56.6 \pm 0.4$ | $61.8 \pm 0.1$ | **$60.2 \pm 0.2$** |
| Distogram | **$60.0 \pm 0.6$** | **$64.2 \pm 0.1$** | $59.2 \pm 0.1$ |

Adding distogram edge features results in consistent improvements across different graph constructions and tasks. In particular, the superior performance of distogram edge features over distance edge features underscores the ability of distograms to capture valuable information regarding RNA conformational variability. These results are particularly interesting given that state-of-the-art structure predictors are generally less accurate for RNA molecules than for proteins (Kretsch et al., 2026), partly owing to the difficulty of tackling RNA flexibility. Please note that these results were obtained without MSA computation, showing that distograms improve performance even without any homology information available.

### 6.4. Robustness to Boltz2 Prediction Quality

In this section, we investigate whether our method is affected by some of the failure modes of Boltz2. We first examine the relationship between the quality of the predicted structures and the performance gains obtained from adding distogram features. We assess prediction quality using the TM-score between the predicted and ground-truth structures. On the RNA tasks, we observe no meaningful correlation between structural prediction quality and the performance gains from distogram features. For example, the improvement in balanced accuracy (resp. MCC) has a Pearson correlation of 0.04 (resp. *−* 0.04) with the TM-score. This suggests that distogram information can improve performance even when the underlying structures are poorly predicted.

We further examine whether the benefits of distogram features depend on MSA depth. Results are reported in Table S7. The gain from distogram features shows no meaningful dependence on MSA depth (all |*r*| *<* 0.08). Although the correlations are slightly negative, their small magnitude does not indicate a systematic relationship. Moreover, the mean F1 gain for the 10% shallowest MSAs is comparable to that observed on the full test set.

## 7. Discussion and conclusions

In this paper, we propose an efficient approach to encode proteins beyond a static structure by using distograms derived from Boltz2, a state-of-the-art protein structure predictor. We propose to enrich graph-based models representing protein structures with edges between residues predicted to be in close proximity according to their corresponding distogram. When the model is compatible, we also include the full distogram distribution as edge features.

For ligand binding site and binding affinity predictions, our approach outperforms existing methods that rely on computationally expensive MD simulations. For the prediction of protein stability, PPI and RNA properties, where MD data were not available, our method also consistently improved performance. Most importantly, across all tasks, the overall best-performing method among different architectures incorporates distograms.

These results highlight the potential of incorporating distogram-derived information to enhance protein as well as RNA representation learning. Nonetheless, our approach presents a few limitations. First, it relies on high-quality distograms, whose accuracy ultimately depends on the strength of coevolutionary signals in the multiple sequence alignment. Second, computing distograms for large proteins or macromolecular complexes can be computationally demanding. Finally, because distograms represent marginalized pairwise distance distributions, they provide an entangled, collective description of conformational variability and may not explicitly capture higher-order, multi-residue dynamic correlations.

An interesting future direction is the application of our method to systems lacking experimental structural information. This setting is particularly challenging, as the performance of structure prediction methods strongly depends on coevolutionary signals and the availability of related structures in the PDB. Consequently, it remains uncertain whether reliable and informative features can be extracted for systems with low-accuracy structural predictions. Nevertheless, our approach represents a promising step toward bridging the gap between the abundance of sequence data and the capabilities of dynamic structural modeling.

## Acknowledgments

This project has received funding from the European Research Council (ERC) under the European Union’s Horizon 2020 research and innovation program (Grant agreement No. 101086685 – bAIes). V.M. is supported by a Junior Springboard Prairie program, funded by the ANR project ANR-23-IACL-0008. W.K. is supported by Fondation pour la Recherche Médicale (FRM) with the following grant number: ECO202406019160. This work was performed using the Maestro cluster at Institut Pasteur and HPC resources from GENCI–IDRIS (Grant AD010315435R1).

## Impact Statement

This paper presents work whose goal is to advance the field of Machine Learning, in particular in the areas of computer aided drug discovery and protein design. There are many potential societal consequences of our work, none which we feel must be specifically highlighted here.

## Appendix

### A. Implementation

The code necessary to run our experiments and reproduce our results can be found at https://anonymous.4open.science/r/DistoDyn-55AB/README.md.

Our distograms were generated using Boltz with default parameters. The MSAs were generated on the fly. For RNA, Boltz does not recommend using MSAs. Alignments were generated with the following command line:

~~~
boltz predict “${INPUT_PATH}” --use_msa_server --model boltz2
     --out_dir “${OUTPUT_DIR}”
~~~

#### A.1 Computational resources

All models were trained using either a single 80GB NVIDIA A100 GPU or a 40GB NVIDIA A40 GPU, depending on memory requirements. In our experiments, the RGAT model required more than 40GB of memory and was therefore always trained on an A100 GPU. The other models were trained on an A40 GPU.

For the binding site prediction tasks, the RGAT model took approximately 20 hours to train, whereas the RGCN and REGNN models required less than 10 hours. The RGAT model was also the slowest on the binding affinity task; however, for this task, all models completed training in under 10 hours. Fine-tuning the thermoMPNN model took 14 hours.

#### A.2 Detailed model definition

##### Relational Graph Convolutional Network (R-GCN)

R-GCN extends the standard GCN by introducing relation-specific weight matrices. Let 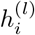 be the representation of node *i* at layer *l*. This representation is modulated by relation-specific weight matrices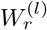, with the convention that *r* = 0 corresponds to self-loops. The message sent by a node *i* across relation *r* is defined as

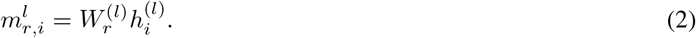

Message passing updates the node representation as:

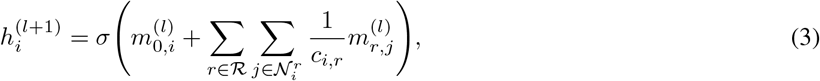

where 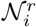 is the set of neighbors of node *i* under relation *r, c*_*i,r*_ is a normalization constant, and *σ* is a non-linear activation function.

##### Relational Graph Attention Network (R-GAT)

R-GAT introduces relation-specific attention coefficients to weight messages from different neighbors:

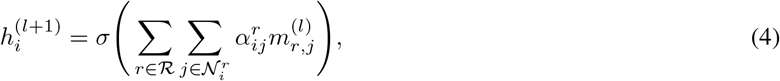

where the attention coefficients 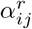 are computed as

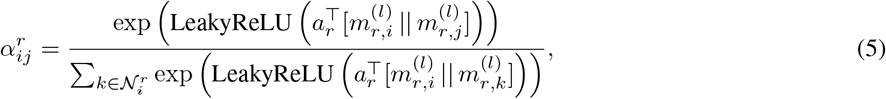

with *a*_*r*_ as a learnable attention vector for relation *r* and || denoting vector concatenation.

##### Relational Equivariant Graph Neural Network (R-EGNN)

To extend EGNN to handle multiple edge types, we create a relational variant (R-EGNN) by processing each relation type separately. Specifically, for each edge type *r* (e.g., distance-based, correlation-based, or distogram-based), a separate EGNN processes the corresponding subgraph, producing relation-specific node and coordinate updates. The outputs from all relations are then combined via summation to obtain the final node embeddings and coordinate updates.

Formally, the node update for relation *r* follows the standard EGNN message passing:

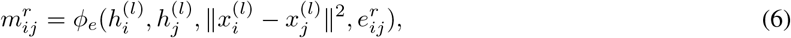

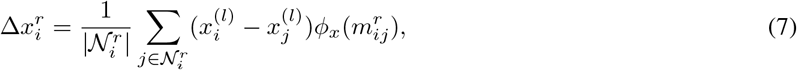

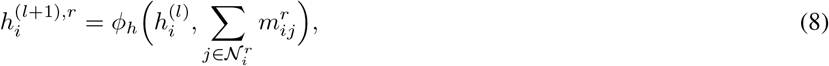

where 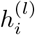 and 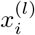 are the feature vector and coordinates of node *i* at layer *l*, 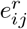is the edge feature vector for relation *r*, and *ϕ*_*e*_, *ϕ*_*x*_, *ϕ*_*h*_ are learnable MLPs.

The final node representation and coordinates are obtained by summing the contributions across all relations:

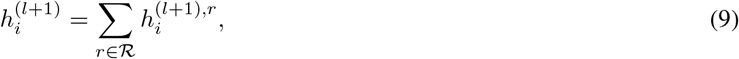

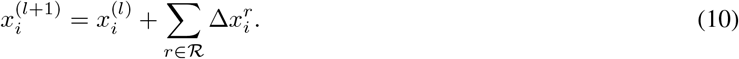

This design allows the model to leverage relation-specific information while maintaining the equivariance properties of EGNN: the predictions are invariant to translations and equivariant to rotations of the input coordinates. By incorporating multiple edge types (distance, correlation, and distogram), R-EGNN captures complementary structural and dynamic information while directly processing 3D node coordinates.

#### A.3 Task-specific details

All models are trained using the AdamW optimizer with learning rate 0.0001, except for ThermoMPNN where different learning rates are used for each module as explained below.

##### A.3.1. Small-molecule binding tasks

It should be noted that our efforts to reproduce the results of Guo et al. (2025) resulted in a systematic performance improvement for Distance-only and Distance+Correlation models, setting a higher bar to beat. However, due to a significant increase in the performance of Distance-only models, adding correlation edges consistently helped to predict binding sites but not affinities, unlike what was previously reported.

###### Data and splits

The MISATO dataset (Siebenmorgen et al., 2024) (https://github.com/t7morgen/ misato-dataset GNU Lesser General Public License v2.1) is used for the binding site and binding affinity prediction task. This dataset contains 19443 protein-ligand complexes extracted from PDBbind (Su et al., 2018; Liu et al., 2017; Wang et al., 2005) (https://www.pdbbind-plus.org.cn/). In the binding site prediction task, the data is partitioned following the split provided with the MISATO dataset. In the binding affinity prediction task, models are trained on the PDBbind 2020 refined set and tested on the core set which contain respectively 5318 and 285 protein–ligand complexes retained after extensive filtering to ensure the quality of both binding affinity data and crystal structures.

###### Graph construction

Following their work, binding sites are composed of protein residues closer than 10 Å to any non-hydrogen atom of the ligand. Entire protein structures are used, resulting in graphs with an average number of nodes equal to 443.

For the binding affinity prediction task, only atoms belonging to the binding pocket are considered, resulting in much smaller graphs (47 residues on average). Moreover, since this task requires representing a ligand, we need to adapt the distogram edge set *ℰ*_*disto*_ introduced in the previous section. Namely, we complement *ℰ*_*disto*_ with atomic radius graph edges on the ligand side, and we connect protein and ligand nodes if one atom of the protein is closer than 8 Å to an atom of the ligand.

###### Model and training setup

For the binding site and binding affinity prediction, we used the models as defined by Guo et al. (2025). More specifically, our models are made up of 5 relational layers (either RGAT or RGCN layers) followed by a 2-layer MLP. For the affinity task, the ligand and protein are pooled separately before being passed to the MLP. We grid-searched dropout rates over the following values : *{*0.0, 0.1, 0.2, 0.3, 0.4*}* for all models.

When using distogram-based edges, the R-GAT and R-EGNN models are also trained with distograms used as edge features. For both tasks, model architectures and hyperparameters are kept identical across different relation combinations, except for the dropout rate, which is tuned to achieve optimal performance independently for each model. A cross-entropy and MSE loss functions are used respectively for the binding-site and affinity prediction task.

We tested rewiring with a varying number of loops (Table S5) and report the best performing setup.

##### A.3.2. Protein stability

Protein stability is measured as the change in free energy Δ*G* between its folded and unfolded states. Upon mutations in the amino acid sequence, stability can be significantly affected. This variation is quantified by ΔΔ*G*, which can be used to compare the effect of different mutations, notably for tasks like protein design.

###### Data and splits

Experiments for the protein stability prediction task are conducted on the mega-scale dataset (Tsuboyama et al., 2023). This dataset contains 776000 high-quality folding stability measurements spanning all single amino acid variants and selected double mutants across 331 natural and 148 de novo–designed protein domains, each 40–72 amino acids long. After removing duplicate and unreliable stability measurements, we end up with 577313 samples. The data are split into training, validation, and test sets following the partitioning scheme used by (Dieckhaus et al., 2024). After removing double mutants, the final dataset contains 298 wild types, accounting for 443,906 protein sequences. Data and splits were held constant across all experiments.

###### Graph construction

ThermoMPNN relies on ProteinMPNN (Dauparas et al., 2022), a popular graph-based structure encoder, to extract structure embeddings. In our experiments, we sought to enrich the ProteinMPNN graphs with distograms. The default number of edges per protein was kept unchanged. Distogram features were passed as additional input to the first linear layer of ProteinMPNN.

###### Model and training setup

The ThermoMPNN model is built around ProteinMPNNDauparas et al. (2022), with an added stability prediction module. This model is a graph neural network (GNN) that uses representations derived from a pretrained sequence-recovery model, ProteinMPNN, to predict the changes in protein thermostability induced by point mutations. The input protein sequence is processed by ProteinMPNN, and the representations produced at its decoder layers are combined with the learned sequence representation to encode the local environment of each residue. These representations are subsequently processed by a lightweight attention (LA) module, which applies self-attention to integrate contextual information and adaptively weight the resulting features. The resulting representation is then fed into a multilayer perceptron (MLP) to estimate the free-energy change, ΔΔ*G*^*°*^, by taking the difference between the predicted Δ*G*^*°*^ values for the mutant and wild-type amino acids.

To train the model, the default MSE loss function was kept. We also include results for SPURS (Li & Luo, 2025), a recent baseline tailored for this task. The reported results are the means *±* standard deviation over five seeds.

For the finetuning of the protein stability tasks, all layers are kept frozen except the initial edge embedding layers, light attention layers and heads. We chose to fine-tune ThermoMPNN instead of training it from scratch. To do so, we kept the ProteinMPNN encoder layers frozen in early epochs and gradually unfroze them. Out of fairness, we also tried to fine-tune their model (allowing for more computations), which ultimately did not affect performance. Different learning rate scheduling are used for these layers. We tried training our models with 6 different learning rate strategies detailed in Table S1 and report our best performance. In addition, the first encoder layer of ProteinMPNN is unfrozen after epoch 10. Then between epoch 10 and 20 the second encoder layer of ProteinMPNN is also unfrozen. Finally, the last encoder layer is unfrozen from epoch 20 until the end of training.

**Table S1.** Setups used for the protein stability prediction task.

| Learning rate setup |  | A | B | C | D | E | F |
| --- | --- | --- | --- | --- | --- | --- | --- |
| Encoder | Initial | $10^{-4}$ | $10^{-4}$ | $10^{-5}$ | $10^{-4}$ | $10^{-3}$ | $10^{-3}$ |
| | Final | $10^{-5}$ | $10^{-5}$ | $10^{-6}$ | $10^{-5}$ | $10^{-4}$ | $10^{-5}$ |
| Edge | Initial | $10^{-3}$ | $10^{-3}$ | $10^{-4}$ | $10^{-4}$ | $10^{-3}$ | $10^{-3}$ |
| | Final | $10^{-5}$ | $10^{-4}$ | $10^{-5}$ | $10^{-5}$ | $10^{-4}$ | $10^{-4}$ |
| Head | Initial | $10^{-3}$ | $10^{-3}$ | $10^{-4}$ | $10^{-4}$ | $10^{-3}$ | $10^{-3}$ |
| | Final | $10^{-5}$ | $10^{-4}$ | $10^{-5}$ | $10^{-5}$ | $10^{-4}$ | $10^{-4}$ |
| Light Attention | Initial | $10^{-3}$ | $10^{-3}$ | $10^{-4}$ | $10^{-4}$ | $10^{-3}$ | $10^{-3}$ |
| | Final | $10^{-5}$ | $10^{-4}$ | $10^{-5}$ | $10^{-5}$ | $10^{-4}$ | $10^{-4}$ |

##### A.3.3. Protein-protein interaction

###### Data and splits

PINDER (Protein INteraction Dataset and Evaluation Resource) is a large-scale benchmark for protein-protein docking. It’s roughly 500x larger than prior state-of-the-art datasets and is the first to pair predicted (AlphaFold2) and apo (unbound) structures with ground-truth complexes, making it useful for training and evaluating flexible docking methods, not just rigid ones. A big part of its contribution is data quality: the curation pipeline specifically filters out problems common in older benchmarks, like structural leakage between train and test clusters, spurious multi-chain interactions, interfacial gaps, and detached chain fragments. For evaluation, PINDER ships curated splits — PINDER-XL is the full test set (1,955 cluster representatives, with 342 apo-paired and 1,747 AF2-paired structures), while PINDER-S is a smaller, diversity-sampled subset (250 clusters, 188 heterodimers + 62 homodimers) for quicker benchmarking. The data used in our experiments covers a wide range of graph sizes (162.5 *±* 70.0 nodes on average), markedly larger than the graphs used in the protein stability task. In total the training, validation and test sets are made up of 450000, 325 and 357 pairs respectively.

###### Data processing

Because all homodimers in this dataset are positive by construction, we restrict evaluation to heterodimers, so as to obtain a balanced and meaningful classification setting. We use MMseqs to remove homodimers from Pinder data. More specifically, the Uniref90 criterion to identify pairs of proteins with high similarity (more than 90% sequence similarity with more than 80% coverage). Similar to (Hu & Ohue, 2025), chains shorter than 35 or longer than 300 amino acids are filtered out, and interactions involving fewer than 8 contact residues are discarded. Negative samples are generated by randomly pairing chains.

###### Model and training setup

Given the structures of two proteins, this task predicts whether they physically interact, using a graph attention network (GAT) applied jointly to both partners. We enrich the SpatialPPIv2 graphs with distogram-based edges, and separately test the addition of distogram distributions as edge features. The four GAT layers used in SpatialPPIv2 are replaced with four RGAT layers to account for the two relation types. The remainder of the architecture is kept unchanged, with the representations of the two chains independently mean-pooled and subsequently concatenated before being processed by a multi-layer perceptron. Similar to (Hu & Ohue, 2025), the binary cross-entropy is used to train the model.

##### A.3.4. Prediction of RNA properties

###### Graph construction

In our RNA experiments, nodes represent residues, and we investigate three possible connectivities: distance-based graphs, 2D+ graphs and 2.5D graphs.

In **distance-based graphs**, each residue is connected to all residues located within a neighborhood of radius 8.0Å (self-loops are removed). To compute distances, we need to pick a representative atom. Following Boltz2, we choose the C2 atom for pyrimidines and the C4 atom for purines. The 8.0Å cutoff was chosen because it results in an average degree of 7.7, which is reasonable given the relatively small size of our graphs. In this setting, we note that there is only one edge type.

In **2D+ graphs**, three distinct edge types are used. A first edge type is created for 5’ to 3’ backbone connections (phosphodiester bonds), a second edge type is added for 3’ to 5’ backbone connections, and a third edge type (bidirectional) for canonical base pairs. These graphs are therefore directed.

In **2.5D graphs**, we additionally model the non-canonical interactions of RNA. 17 edge types are introduced in addition to the three edge types of 2D+ graphs, corresponding to each of the base pair geometries as defined by the Leontis-Westhof nomenclature of base pairs (Leontis & Westhof, 1998). These edges were shown to improve the performance of machine learning approaches for drug design applications (Oliver et al., 2020), and on a recent benchmark for RNA 3D structure-function modeling (Wyss et al., 2025).

For each setting, we run experiments without any edge features, with distogram-based edge features, and with distance-based edge features, where distance is encoded with 16 radial basis function.

###### Model and training setup

The experiments were carried out using relational graph attention networks (RGAT). This architecture was chosen for its ability to natively handle simultaneously distinct edge types and continuous edge features. For a fair comparison across settings, we also used the RGAT architecture for the experiments carried out without distance- or distogram-based edge features. The implementation was performed using RNAglib Tasks (Wyss et al., 2025).The loss function is weighted by the relative occurrences of the classes in our datasets. The hyperparameters chosen are reported below:

**Table S2.** Hyperparameter settings used for the experimental evaluation on RNA structures.

| Hyperparameter | Value |
| --- | --- |
| Number of Layers | 3 |
| Hidden Channels | 128 |
| Training Epochs | 200 |
| Batch Size | 8 |
| Dropout Rate | 0.5 |
| Loss Weights | Ratio-based |
| Learning Rate | $\{10^{-3}, 10^{-4}\}$ (Tuned via validation) |

### B. Supplementary results

#### B.1. Extended results on datasets where MD is available

The results with other models is presented in Table S3.

#### B.2. Complementarity to Correlation

We have conducted two additional experiments to investigate the synergies between using MD-derived correlation and distogram. The first one uses the MD data to compute an empirical distogram, by averaging measurements of the pairwise distances over conformations. This allows to use exactly the same setup as the one we use when constructing distogram-based edges, ensuring direct comparison.

Our second experiment looks at the performance of a model trained with both distogram-edges and correlation edges (in the classic setup, not the empirical distogram from previous experiment). This is useful to explore if complementary information is contained in the two modalities. Results are presented in Table S4.

First, we see that the performance of the MD-derived distogram is generally close to the one of correlation edges (with reduced accuracy and enhance recall), and far from the performance of our approach. This indicates that the performance gap between distogram and correlation edges does not stem from the specific encoding of the MD information, but rather that the distogram is richer in information.

Second, we see that incorporating correlation edges in addition to distance and distogram edges yields little to no improvement in either binding site or binding affinity prediction, suggesting that the correlation information is largely captured by the existing distance and distogram relationships, since both capture related aspects of the system’s dynamic behavior.

**Table S3.** Results on the binding site prediction task (Top, average of 443 nodes) and binding affinity prediction task (Bottom, 47 nodes on average) for the R-EGNN model. We compare various dynamic-encoding approaches using *ℰ*_*dist*_ alone or combined with *ℰ*_*corr*_ or *ℰ*_*disto*_. Distogram-based edge features are also incorporated where compatible. Best-performing models are shown in **bold**, and second-best are underlined.

| Binding Site Prediction (mean nodes = 443) |  |  |  |  |  |
| --- | --- | --- | --- | --- | --- |
| Model | Graph Type | Acc. (↑) | Prec. (↑) | Rec. (↑) | F1 (↑) |
| R-GCN | Distance | 0.761 | 0.187 | <u>0.420</u> | 0.259 |
|  | + Correlation | **0.809** | <u>0.217</u> | 0.357 | <u>0.270</u> |
|  | + Distogram | <u>0.782</u> | **0.236** | **0.535** | **0.328** |
| R-GAT | Distance | 0.753 | 0.196 | 0.478 | 0.278 |
|  | + Correlation | 0.791 | 0.229 | 0.469 | 0.308 |
|  | + Distogram | <u>0.814</u> | <u>0.289</u> | <u>0.598</u> | <u>0.390</u> |
|  | + Distogram + Features | **0.816** | **0.298** | **0.627** | **0.404** |
| R-EGNN | Distance | 0.832 | 0.282 | 0.444 | 0.345 |
|  | + Correlation | **0.882** | **0.393** | 0.350 | 0.370 |
|  | + Distogram | <u>0.861</u> | <u>0.376</u> | **0.606** | **0.464** |
|  | + Distogram + Features | 0.859 | 0.371 | <u>0.602</u> | <u>0.459</u> |
| Binding Affinity Prediction (mean nodes = 47) |  |  |  |  |  |
| Model | Graph Type | MAE (↓) | RMSE (↓) | Pearson (↑) | Spearman (↑) |
| R-GCN | Distance | <u>1.244</u> | <u>1.562</u> | <u>0.689</u> | <u>0.656</u> |
|  | + Correlation | 1.299 | 1.601 | 0.673 | 0.637 |
|  | + Distogram | **1.195** | **1.487** | **0.731** | **0.694** |
| R-GAT | Distance | 1.226 | 1.562 | 0.691 | 0.658 |
|  | + Correlation | 1.280 | 1.568 | 0.686 | 0.656 |
|  | + Distogram | **1.176** | **1.487** | **0.740** | **0.713** |
|  | + Distogram + Features | <u>1.179</u> | <u>1.492</u> | <u>0.722</u> | <u>0.693</u> |
| R-EGNN | Distance | 1.296 | 1.623 | 0.666 | 0.642 |
|  | + Correlation | 1.357 | 1.713 | 0.611 | 0.576 |
|  | + Distogram | <u>1.275</u> | <u>1.560</u> | <u>0.699</u> | <u>0.674</u> |
|  | + Distogram + Features | **1.208** | **1.479** | **0.736** | **0.725** |

**Table S4.** Other ways to incorporate MD-correlation information.

| Model | Graph Type | Accuracy (↑) | Precision (↑) | Recall (↑) | F1 score (↑) |
| --- | --- | --- | --- | --- | --- |
| R-GCN | Distance | 0.761 | 0.187 | 0.42 | 0.259 |
|  | Distance + Correlation | **0.809** | 0.217 | 0.357 | 0.270 |
|  | Distance + MD Distogram | 0.759 | 0.184 | 0.415 | 0.255 |
|  | Distance + Distogram | 0.782 | <u>0.236</u> | **0.535** | **0.328** |
|  | Distance + Distogram + Correlation | <u>0.803</u> | **0.246** | <u>0.474</u> | <u>0.324</u> |
| R-GAT | Distance | 0.753 | 0.196 | 0.478 | 0.278 |
|  | Distance + Correlation | 0.791 | 0.229 | 0.469 | 0.308 |
|  | Distance + MD Distogram | 0.746 | 0.197 | 0.504 | 0.283 |
|  | Distance + Distogram | 0.814 | 0.289 | <u>0.598</u> | 0.39 |
|  | Distance + Distogram + Features | <u>0.816</u> | **0.298** | **0.627** | **0.404** |
|  | Distance + Distogram + Correlation | **0.819** | <u>0.293</u> | 0.584 | <u>0.391</u> |
| R-EGNN | Distance | 0.832 | 0.282 | 0.444 | 0.345 |
|  | Distance + Correlation | <u>0.882</u> | <u>0.393</u> | 0.35 | 0.37 |
|  | Distance + MD Distogram | 0.835 | 0.305 | 0.516 | 0.383 |
|  | Distance + Distogram | 0.861 | 0.376 | <u>0.606</u> | <u>0.464</u> |
|  | Distance + Distogram + Features | 0.859 | 0.371 | 0.602 | 0.459 |
|  | Distance + Distogram + Correlation | **0.887** | **0.438** | **0.496** | **0.465** |

#### B.3. Additional rewiring results

We tested various values of rewiring for the RGCN on the binding site task, and report results in Table S5. Performance increases with the number of steps up until 5000, and then decreases. For all values, pure rewiring underperforms distogram edges. Moreover, in all cases, adding distogram information enhances performance.

**Table S5.** Rewiring results for different numbers of rewiring steps.

| Graph Type | Num Rewire | Accuracy (↑) | Precision (↑) | Recall (↑) | F1 score (↑) |
| --- | --- | --- | --- | --- | --- |
| Distance | - | 0.761 | 0.187 | 0.42 | 0.259 |
| Distance + Distogram | - | 0.782 | 0.236 | 0.535 | 0.328 |
| Distance + Rewired | 10 | 0.776 | 0.193 | 0.394 | 0.259 |
|  | 100 | 0.769 | 0.19 | 0.405 | 0.258 |
|  | 1000 | 0.783 | 0.224 | 0.479 | 0.305 |
|  | 5000 | 0.784 | 0.228 | 0.491 | 0.311 |
|  | 10000 | 0.769 | 0.216 | 0.504 | 0.303 |
| Distance + Rewired + Distogram | 100 | 0.78 | 0.239 | 0.554 | 0.334 |
|  | 1000 | 0.778 | 0.241 | 0.574 | 0.339 |
|  | 5000 | <u>0.79</u> | <u>0.253</u> | **0.568** | <u>0.35</u> |
|  | 10000 | **0.8** | **0.26** | <u>0.548</u> | **0.352** |

We expand our analysis to the binding affinity task and to the different models used. We repeated the exploration of various number of steps. **We always report results corresponding to the number of steps with the strongest performance on the model without distograms**. Our results are presented in Table S6.

On the binding site prediction task, rewiring consistently helps and the rewiring and distogram effects are consistently synergistic. For the RGCN and RGAT models, rewiring strongly helps and the best performances are obtained with by combining rewiring and distograms. For the REGNN model, rewiring only moderately enhances performance compared to distance-based edges, so the combined model has a similar performance to the distogram-only model.

**Table S6.** Extensive rewiring results.

| Model | Graph Type | Accuracy (↑) | Precision (↑) | Recall (↑) | F1 score (↑) |
| --- | --- | --- | --- | --- | --- |
| R-GCN | Distance | 0.761 | 0.187 | 0.42 | 0.259 |
|  | Distance + Distogram | 0.782 | <u>0.236</u> | <u>0.535</u> | <u>0.328</u> |
|  | Distance + Random Edges | 0.772 | 0.191 | 0.401 | 0.259 |
|  | Distance_rewired_5000 | <u>0.784</u> | 0.228 | 0.491 | 0.311 |
|  | Distance_rewired_5000 + distogram | **0.79** | **0.253** | **0.568** | **0.35** |
| R-GAT | Distance | 0.753 | 0.196 | 0.478 | 0.278 |
|  | Distance + Distogram | 0.814 | 0.289 | <u>0.598</u> | 0.39 |
|  | Distance + Random Edges | **0.825** | <u>0.298</u> | 0.567 | <u>0.391</u> |
|  | Distance_rewired_5000 | 0.798 | 0.248 | 0.509 | 0.334 |
|  | distance_rewired_5000 + distogram | <u>0.823</u> | **0.306** | **0.614** | **0.408** |
| R-EGNN | Distance | 0.832 | 0.282 | 0.444 | 0.345 |
|  | Distance + Distogram | **0.861** | **0.376** | <u>0.606</u> | **0.464** |
|  | Distance + Random Edges | 0.849 | 0.335 | 0.526 | 0.409 |
|  | Distance_rewired_100 | 0.837 | 0.306 | 0.504 | 0.381 |
|  | distance_rewired_100 + distogram | <u>0.859</u> | <u>0.372</u> | **0.608** | <u>0.462</u> |

| Model | Graph Type | MAE (↓) | RMSE (↓) | Pearson R (↑) | Spearman R (↑) |
| --- | --- | --- | --- | --- | --- |
| R-GCN | Distance | 1.244 | 1.562 | 0.689 | 0.656 |
|  | Distance + Distogram | <u>1.195</u> | <u>1.487</u> | <u>0.731</u> | <u>0.694</u> |
|  | Distance + Random Edges | 1.368 | 1.715 | 0.649 | 0.619 |
|  | Distance_rewired_10 | 1.265 | 1.583 | 0.679 | 0.64 |
|  | Distance_rewired_10 + distogram | **1.159** | **1.457** | **0.737** | **0.707** |
| R-GAT | Distance | 1.226 | 1.562 | 0.691 | 0.658 |
|  | Distance + Distogram | **1.176** | **1.487** | **0.74** | **0.713** |
|  | Distance + Random Edges | 1.205 | <u>1.491</u> | <u>0.73</u> | <u>0.704</u> |
|  | Distance_rewired_5000 | 1.243 | 1.559 | 0.7 | 0.687 |
|  | Distance_rewired_5000 + distogram | 1.21 | 1.527 | 0.717 | 0.699 |
|  | Distance + Distogram + Features | <u>1.179</u> | <u>1.492</u> | 0.722 | 0.693 |
| R-EGNN | Distance | 1.296 | 1.623 | 0.666 | 0.642 |
|  | Distance + Distogram | 1.275 | 1.56 | 0.699 | 0.674 |
|  | Distance + Random Edges | 1.286 | 1.602 | <u>0.712</u> | <u>0.688</u> |
|  | Distance_rewired_1000 | <u>1.267</u> | <u>1.552</u> | 0.7 | 0.683 |
|  | Distance_rewired_1000 + distogram | 1.314 | 1.616 | 0.673 | 0.631 |
|  | Distance + Distogram + Features | **1.208** | **1.479** | **0.736** | **0.725** |

On the binding affinity task, rewiring does not consistently increase the performance of distance-based graphs. We attribute this effect to the moderate size of the graphs, as well as to their heterogeneous nature where nodes represent either protein residues or ligand atoms. Interestingly, even in scenarios where rewiring decreases performance, it can help when used in conjunction with distograms (RGCN performance).

*Overall, the best performing method involves distograms across all models, metrics and tasks*.

#### B.4. Correlation with MSA depth

We next test whether the gains from distogram features vary with MSA depth. For each protein system in the MISATO binding-site task (*n* = 1610), we measure the performance gain from adding distogram features and correlate it with the MSA depth used by Boltz2. We additionally compare the gains observed for the 10% of systems with the shallowest MSAs with those obtained on the full test set (Table S7). Across all models, the correlation between performance gain and MSA depth is negligible (all |*r*| *<* 0.08); the correlations are weakly negative, but too small in magnitude to suggest any systematic relationship. Consistent with this, the mean F1 gain on the 10% of systems with the shallowest MSAs is comparable to the gain observed across the full test set.

**Table S7.** Relationship between MSA depth and model performance gains.

| Model | Pearson $r$ (gain vs. depth) | Mean F1 Gain | |
| --- | --- | --- | --- |
|  |  | Whole Test | Shallowest 10% |
| RGCN | -0.058 | +0.071 | +0.099 |
| RGAT | -0.012 | +0.068 | +0.038 |
| REGNN | -0.066 | +0.065 | +0.063 |

## Notes

### Competing Interest Statement

The authors have declared no competing interest.

### Summary of Updates

Add PPI benchmark + improved analysis

